# CREWdb 1.0: Optimizing Chromatin Readers, Erasers, and Writers Database using Machine Learning-Based Approach

**DOI:** 10.1101/2022.06.02.494594

**Authors:** Maya Natesan, Reetika Ghag, Mitchell Kong, Min Shi, Shamim Mollah

**Affiliations:** Institute for Informatics, Washington University School of Medicine in St. Louis, 600 Euclid Ave. MO 63110; Department of Bioengineering, University of California, San Diego, 9500 Gilman Drive, La Jolla, CA 92093; Department of Genetics, Washington University School of Medicine in St. Louis, 600 Euclid Ave. MO 63110

## Abstract

Aberration in heterochromatin and euchromatin states contributes to various disease phenotypes. The transcriptional regulation between these two states is significantly governed by post-translational modifications made by three functional types of chromatin regulators: readers, writers, and erasers. Writers introduce a chemical modification to DNA and histone tails, readers bind the modification to histone tails using specialized domains, and erasers remove the modification introduced by writers. Altered regulation of these chromatin regulators results in complex diseases such as cancer, neurodevelopmental diseases, myocardial diseases, kidney diseases, and embryonic development. Due to the reversible nature of chromatin modifications, we can develop therapeutic approaches targeting these chromatin regulators. However, a limited number of chromatin regulators have been identified thus far, and a subset of them are ambiguously classified as multiple chromatin regulator functional types. Thus, we have developed machine learning-based approaches to predict and classify the functional roles of chromatin regulator proteins, thereby optimizing the accuracy of the first comprehensive database of chromatin regulators known as *CREWdb*.

**GitHub URL:** CREWdb source code is available at https://github.com/smollahlab/CREWdb

**Database URL:** CREWdb webtool is available at http://mollahlab.wustl.edu/crewdb

## Introduction

DNA serves as the biological basis for cellular replication, transcription, and translation—processes that govern gene expression in cells. While cells contain identical genetic material, their ability to maintain differential gene expression and unique functionality is due mainly to the regulation of their DNA by epigenetic modifiers. Epigenetic modifications performed by catalyzing enzymes are key players in gene expression patterns during cell differentiation and development (1).

In eukaryotic cells, DNA wraps around proteins known as histones, forming condensed coils. Epigenetic modifiers act on these newly translated histone proteins, influencing chromatin accessibility and gene expression (2). There are two main types of chromatin remodeling: histone modifications and ATP-dependent chromatin remodeling complexes. The former is a key driver in many known diseases, such as acute myeloid leukemia, breast cancer, and neurodegenerative disorders (3). These histone modifications cause changes in DNA accessibility by neutralizing or introducing charges on chromatin, thereby weakening or strengthening nucleosome interactions between histone groups (4). Stronger interactions condense the DNA and result in transcriptional repression, while weaker interactions uncoil the DNA and result in transcriptional activation.

The enzymes responsible for the condensation and uncoiling of chromatin are chromatin regulators, characterized as readers, erasers, or writers (REW). Writers introduce functional groups to DNA and histones, including those under histone methyltransferase and histone acetyltransferase domains (5). Readers contain specialized domains that recognize and interpret these functional groups and include proteins that contain methyl-lysine-recognition motifs such as bromodomains and chromodomains (6). Erasers remove the chemical modifications introduced by writers and include those under histone demethylase and histone deacetylase domains. As epigenetic drivers, these chromatin regulators play critical roles in regulating gene expression. Therefore, a comprehensive understanding of chromatin regulators is crucial for further research and development in treatment therapies for many diseases, including cancers and genetic disorders.

However, current efforts to compile descriptors of epigenetic regulators are limited. Many studies focus on specific markers and enzymes (7) or tissue-specific cancers (6) rather than all chromatin regulators across all diseases and subtypes. In addition, the available databases that aim to address this problem are currently incomplete, lack coverage, and are inconsistent (8). Thus, our database, the Chromatin Readers, Erasers, and Writers database (CREWdb), aims to mitigate this problem by providing a comprehensive collection of bonafide chromatin regulators and their molecular features across multiple diseases. We initially curated these regulators using various known sources from publicly available databases and published studies, including Chromatin Regulator to Cancer (CR2Cancer) (9), Writers, Erasers, Readers of Acetylation and Methylation proteins database (WERAM DB) (10), Gillette (6), Nicholson (11), and Hyun (2). We developed a web interface where researchers can access and gain more information about the functional roles and attributes of their chromatin regulator protein/s of interest. In this study, we aim to optimize the CREWdb by developing a machine-learning pipeline to predict and classify proteins as readers, writers, and erasers using disease-specific features. We focused on predicting chromatin regulator roles in cancer as a starting point.

This paper will 1) discuss the preliminary development of CREWdb and the web interface, 2) outline the proposed machine learning pipeline, including data acquisition and preprocessing, feature selection, training, and testing, and 3) demonstrate the machine learning component of the chromatin regulator classification model for the CREWdb.

## Material and methods

We adopted a well-established data pre-processing and machine learning pipeline to optimize the CREWdb. We integrated curated datasets, performed feature enrichment through feature selection and class rebalancing techniques, and implemented machine learning models to classify chromatin remodeler proteins into their functional roles as readers, writers, or erasers.

### CREWdb database overview

#### Preliminary Development of CREWdb

Preliminary processing for the CREWdb was first performed on several sets of data obtained from established literature sources, including publicly available databases and published research studies. The databases include CR2Cancer, WERAM DB, Cancer Cell Line Encyclopedia (CCLE), and The Cancer Genome Atlas (TCGA), and the studies include Nicholson, Hyun, and Gillette—all of which contain gene names, functional classifications, and validated chromatin remodeler functions as reader, writer, and eraser designations. A mapping script was generated to match CREW characterizations to corresponding genes, resulting in 1484 overall recorded CREW-validated proteins. A brief description of CREWdb construction and each data source is described below.

#### System flow of CREWdb and Web Interface

We developed the CREWdb system by employing a three-tier architecture (12) which includes a data layer, application logic layer, and user interface, as shown in Figure 1. The data layer involves storing the curated data in a MySQL relational database where relationships are represented as tables and defined using primary and foreign key constraints. These tables are normalized to reduce data redundancy and follow relational databases’ ACID (13) properties. Using PHP scripting language, the application layer is built to query the MySQL database, retrieve data, and apply processing logic. Using Apache Webserver, we then constructed a web application. The client-side web interface allows users to interact with the web application through a request-response model (14). The user requests from the client side are sent to the application hosted on the server, and the results generated by the application layer from the data tier are relayed back to the web interface using server responses (Figure 1).

**Figure 1:**
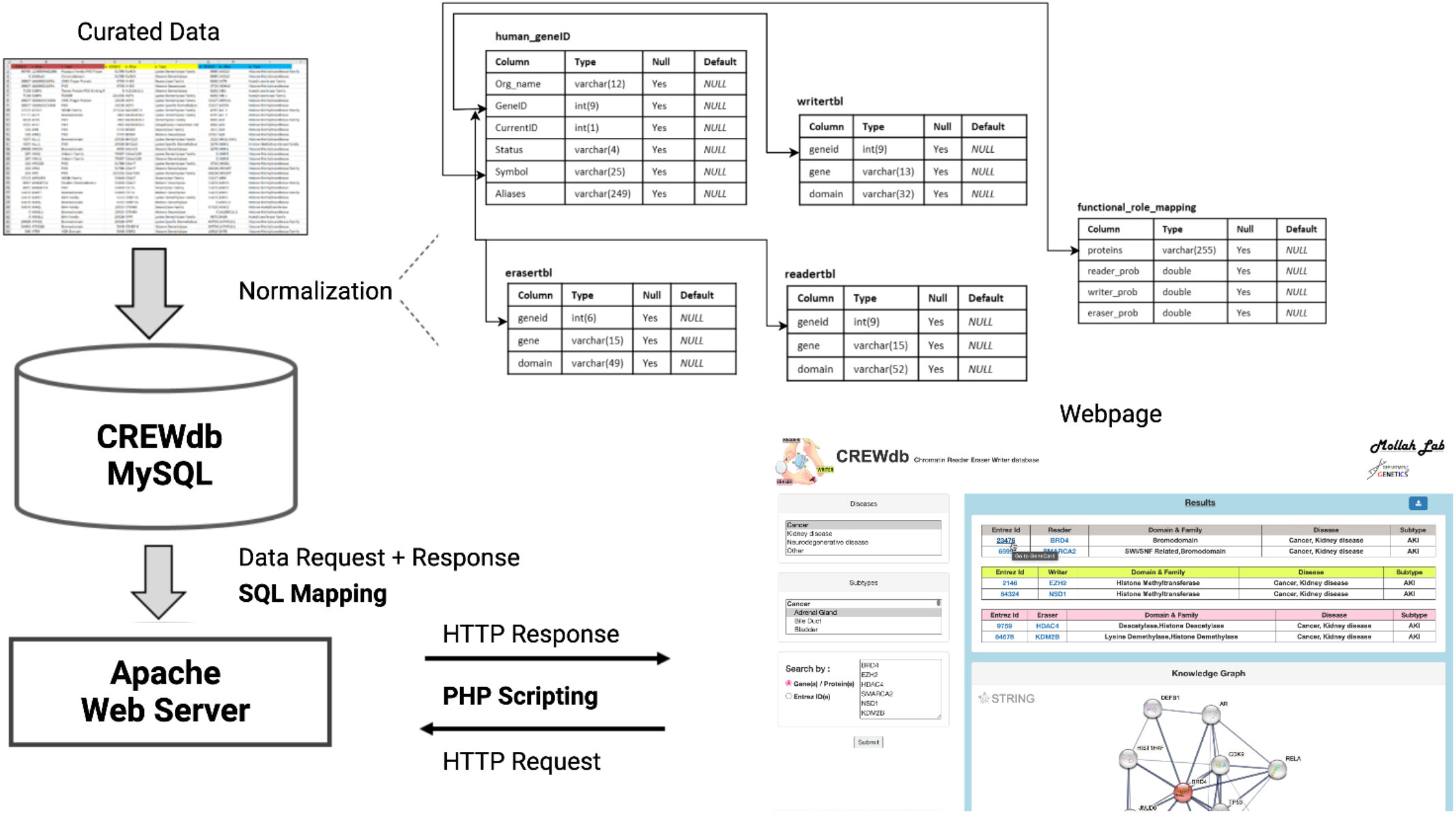
CREWdb architecture diagram and System Flow. The system flow represents the components corresponding to data, logic layers and the user interface of the CREWdb system and their interactions.

### Dataset acquisition

#### CR2Cancer

CR2Cancer is a database released in 2017 and contains genomic, transcriptomic, proteomic, clinical, and functional information on over 400 chromatin regulators across various cancer types [9]. A list of 429 chromatin regulators was generated from three papers (7,8,15). Among these 429 chromatin regulators, 167 were mapped to cancer genes in the Oncogene database (ONGene) (16), Tumor suppressor Gene Database (TSGene) (17,18), and Network of Cancer Genes (NCG) (19) databases. The full name, aliases, chromosome location, and external IDs were annotated for each regulator. In addition, somatic mutation, RNA-seq, DNA copy number, methylation, Reverse Protein Phase Array (RPPA), and clinical data were obtained from TCGA (7); the site-specific substrates and products were compiled from EpiFactors (20); mutation profiles and gene expression microarray of cancer cell lines from CCLE; and the RNA-Seg and antibody staining data of healthy human tissues from HPA (21). For each gene, CR2Cancer provides the (i) functional category, including protein domain, description, substrates, and products of chromatin regulator-specific reactions, (ii) mutation category, including the mutational landscape of primary tumor tissues, (iii) modification category, including post-translational modifications from UniProt, (iv) RNA expression, (v) correlational analysis between RNA expression and semantic copy number alteration, (vi) correlational analysis between RNA expression and promoter methylation level (vii) clinical category, (viii) proteomic, (ix) target category, (x) drug and gene-centric networks, (xi) and protein-protein interaction as well as other interactions derived from literature sources.

#### CCLE

The Cancer Cell Line Encyclopedia (CCLE) database provides genetic characterizations of human cancer cell lines, providing detailed genomic analyses and visualizations across multiple genes and corresponding cancer cell lines (22). The CCLE contains sequencing data aligned to the hg19 broad variant reference genome for nearly 1000 cancer cell lines, including RNA seq, Whole Genome Seq, and Whole Exome Seq.

#### WERAM DB

The Writers, Erasers, Readers of Acetylation and Methylation (WERAM) proteins database provides 584 non-redundant protein data of 8 organisms, including *H. sapiens, M. musculus, R. norvegicus, D. melanogaster, C. elegans, A. thaliana, S. pombe*, and *S. cerevisiae*. This data is further classified into 15 families for histone acetylation writers, erasers, and readers and 32 families for histone methylation writers, erasers, and readers, respectively (10).

The 584 non-redundant protein data consist of histone regulators that were obtained from scientific literature published after 2011. These papers were selected through PubMed Advanced Search Builder, and specific filters were applied for histone acetylation and histone methylation. Protein domain and functional domain information for the extracted histone regulators were obtained from annotations in the UniProt and Pfam databases, respectively.

### Machine Learning pipeline

#### Preprocessing and organization

In order to optimize the preliminary CREWdb described above, we collected additional data on biologically relevant features and developed a machine learning pipeline to predict chromatin regulator functionality as shown in Figure 2.

**Figure 2:**
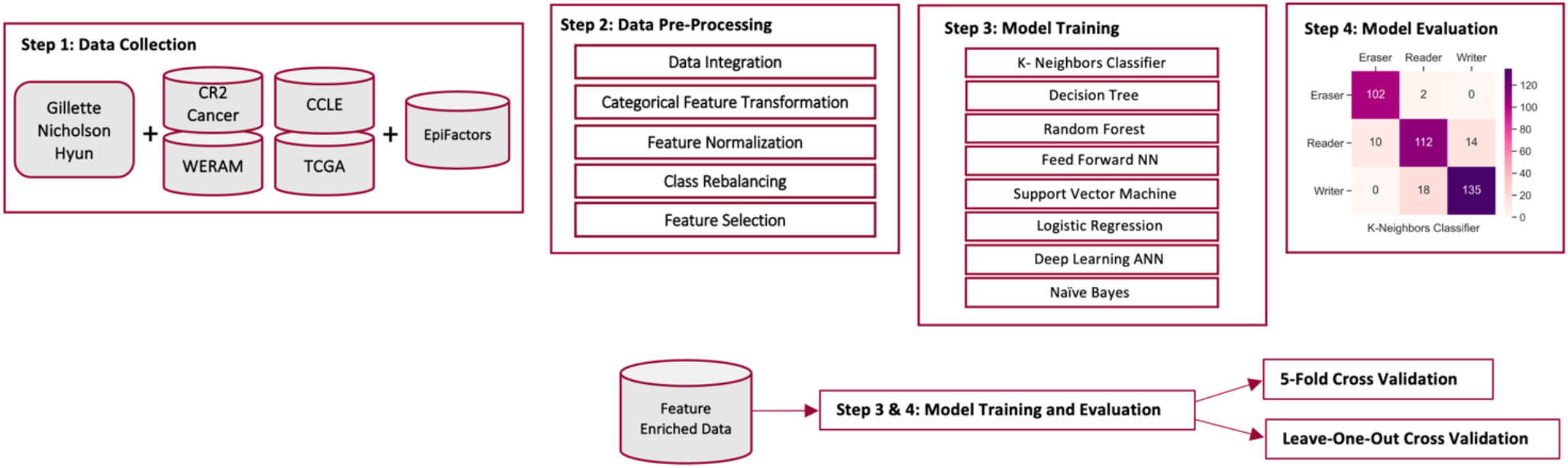
The overview of machine learning pipeline. Schematic representation of data collection, pre-processing, model training, and model evaluation.

Additional data for the machine learning pipeline was extracted from the EpiFactors database. The data curation consisted of 11 unique epigenetic features for 721 proteins. These 11 selected features included gene/protein symbol, HGNC approved name, Pfam domain, HCNC gene family tag, HCNC gene family designation, functional classification, associated histone modification, associated protein complex, target molecule, target entity, and product. A brief description of the EpiFactors database is described below.

#### EpiFactors

The EpiFactors database contains information on epigenetic regulators, their complexes, targets, and products across 458 human primary cell samples, for 815 proteins, including 95 histones and protamines (20). The database includes genes that are specifically defined as epigenetic factors by the founders and include proteins that act as histones, histone variants, or protamines, perform post-translational modifications or can recognize those modifications, change the general structure of chromatin, assist histone folding and assembly, act upon modifications of DNA or RNA, or protein cofactors forming complexes with epigenetic factors (20).

The data for Epifactors database is curated from multiple sources including The Histone Infobase (HIstome) (23) and relevant research papers and reviews. Histones, protamine, their modifying enzymes, DNA methylation enzymes, and chromatin-remodeling proteins were obtained from UniProt through keyword searches. Information on protein complexes is manually extracted from the ‘function’ field in Uniprot. Each protein documented in Epifactors database is a bonified epigenetic regulator validated by peer reviewed published literature (20).

#### Data Integration

Of the 721 proteins extracted from the EpiFactors database, 397 proteins were matched to our CREW characterizations of the initial 1484 CREW-validated proteins. The 11 epigenetic features were then integrated for these 397 proteins, forming the primary dataset for the regulator classification model.

#### Data Transformation

Preliminary data pre-processing included further filtering of the 397 proteins with missing features, nominal encoding and feature normalization that resulted in 393 proteins. Categorical feature transformation was then performed by mapping categorical values such as domain names and family tags to numerical values using one-hot encoding (24) from the Sklearn preprocessing package (25).

#### Class Distribution Rebalancing

The class distribution of the 393 proteins/samples, readers (136 samples), writers (153 samples), and erasers (104 samples) were imbalanced in the dataset as shown in Figure 3A. To mitigate the risk of inaccurate predictions, we then performed a class distribution rebalancing process using a synthetic minority over-sampling technique (SMOTE) (26). SMOTE is an over-sampling method in which the minority class is over-sampled by creating “synthetic” samples. By taking each minority class sample and introducing synthetic samples along the line segments joining any or all minority class nearest neighbors, the number of samples in the minority class increases to match the number of samples in the majority class. For the reader and eraser categories, 17 and 49 samples were generated, respectively, to match the number of samples in the writer’s category.

**Figure 3:**
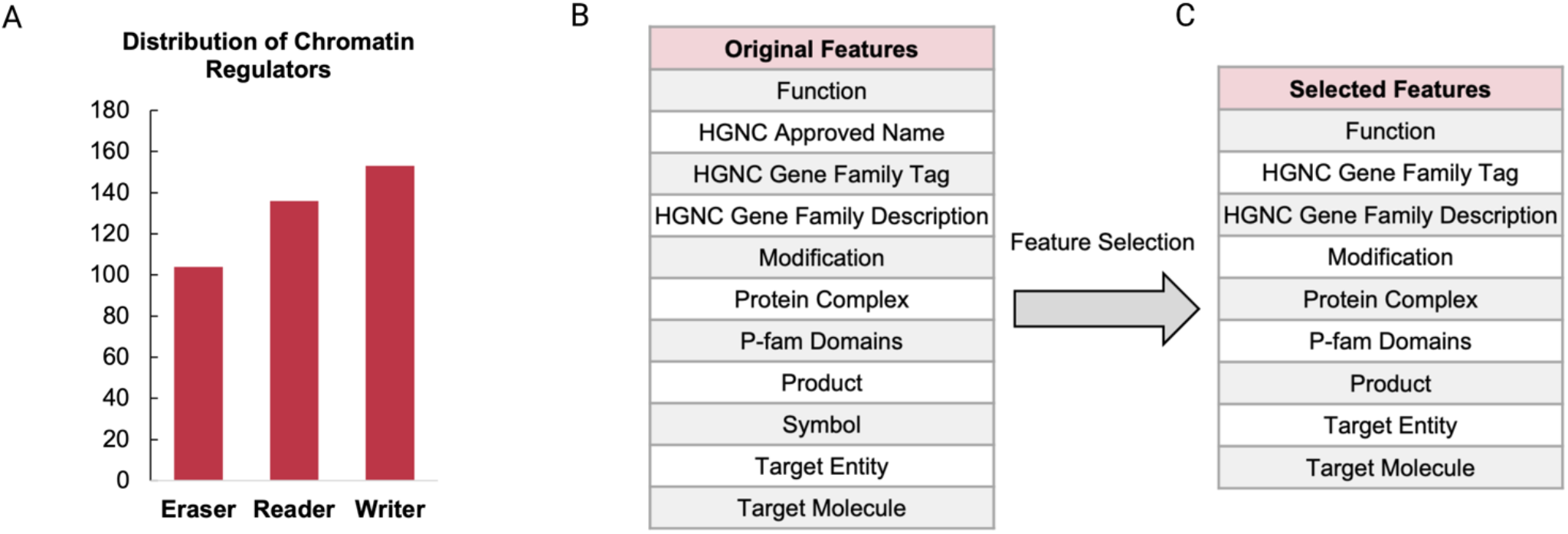
(A) The distribution of readers, writers, and erasers given the collected data. (B) Features listed before and (C) after feature selection

#### Feature Selection

In the final pre-processing step, feature selection was performed on the 11 features (Figure 3B) to select a subset of highly confident features. We adopted a ANOVA based feature selection technique (27) to compute impurity-based feature importance at a threshold of 0.07. With this feature selection technique, 2 features were removed. The final dataset contained 393 proteins with 9 highly confident features shown in Figure 3C.

#### Training and Testing

An agnostic approach was adopted for the machine learning pipeline. We chose 8 different machine models for prediction: Decision Tree, Support Vector Machine, Random Forest, Feed Forward Neural Network, K-Nearest Neighbors, Logistic Regression, Naive Bayes and Deep Neural Network. We used 5-fold and leave-one-out cross validations for training and testing. These 8 prediction models are used for training and testing, as described below.

#### The Prediction Models

- **K-Nearest Neighbor Classifier** (28): A non-parametric supervised learning method that stores instances of training data such that a distance metric such as Euclidean distance is calculated between the test data and each instance of training data. The calculated distances are then sorted in ascending order based on their values and then the top number of *k*, the number of nearest neighbors chosen, rows are extracted and the most frequent class for each row is returned as the predicted class.
- **Decision Tree** (29): A non-parametric supervised learning method that aims to predict the value of a target variable by learning simple decision rules inferred from the data features. The model is obtained by recursively partitioning the data space and fitting a simple prediction model within each partition, where the partitioning can be represented graphically as a decision tree. Entropy is used to measure the quality of a split.
- **Random Forest** (30): A meta-estimator model consisting of a collection of tree-structured (decision tree) classifiers on various sub-samples of the dataset. After a large number of trees is generated, each tree casts a unit vote for determining the most popular class for a given new input sample.
- **Feed Forward Neural Networks (FNN)** (31): An artificial neural network model consisting of multi-layer perceptron (e.g., fully connected). It takes a feature vector as input and outputs a probability value to indicate which class to be assigned to for the input data. FNN optimizes the log-loss function using stochastic gradient descent.
- **Support Vector Machine (SVM)** (32): A supervised learning machine for two-group classification problems. The main idea for SVM is to map the input feature vectors into some high-dimensional feature space Z through some non-linear mapping kernel function. Then, in this space, a linear decision surface (or hyperplane) is constructed with special properties that ensure the high generalization ability of the model to new data.
- **Logistic Regression** (33–35): A statistical model that estimates the probability of each data point classification by using a linear combination of observed features and problem-specific parameters. The values of the parameter are derived from the training data, which are then used to make the predictions on the testing data.
- **Deep Neural Network (Deep Learning ANN)** (36,37): A deep neural network (DNN) is an artificial neural network (ANN) with multiple hidden layers between the input and the output layers. DNNs can capture non-linear patterns where every layer captures a primitive pattern. The DNN uses two hidden layers in addition to the input and the output layer, which outputs the probability of each of the class variables. The weights learned by the model are adjusted using RELU activation function and backpropagation algorithm such as Adam optimizer (38).
- **Naïve Bayes** (39,40): A supervised machine learning algorithm based on the Bayes theorem. The naïve assumption of Bayes theorem assumes conditional independence among all the feature variables to find the value of the class variable. The probability of the class variable value can be estimated using maximum a posteriori (MAP) estimation. The Gaussian Naïve Bayes variant, assumes that each of the class (outcome) variables follow a Gaussian distribution and outputs the probability of each of the class variables for all given features. It requires a small number of training data to calculate necessary parameters. The prediction is based on the following objective function:

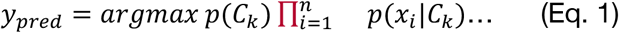

## Results and Evaluation

We evaluated 8 models using 5-fold and leave-one-out cross validation (41) and compared their prediction results by calculating prediction accuracies (Eq. 2), F1 scores (Eq. 5), and AUC (42,43) scores shown in Figure 4. The prediction accuracies and AUC scores were generated through the respective Sklearn metrics for each model. The precision and recall metrics were calculated using Eqs. 3 and 4 and were subsequently used to calculate the F1 scores using Eq. 5.

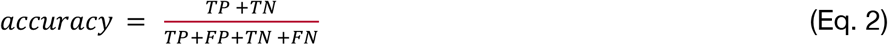

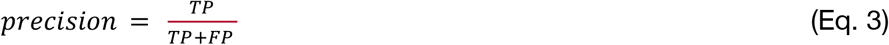

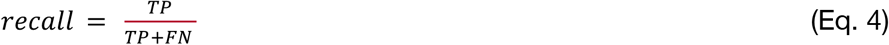

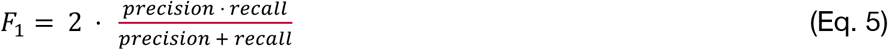

**Figure 4:**
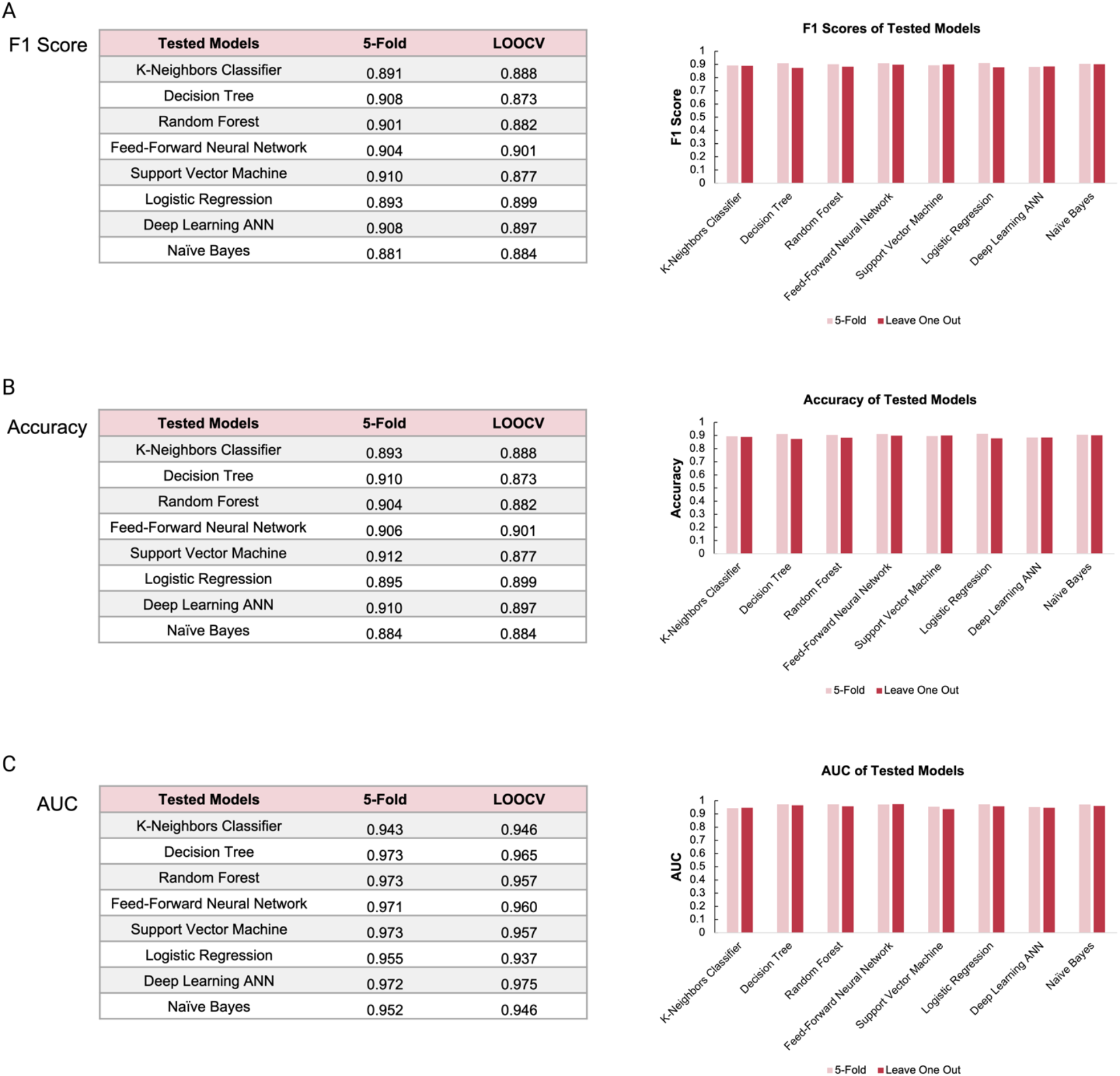
Evaluation metrics for each prediction model. (A) F1-Scores (B) Accuracies (C) AUC values listed and plotted for each classification model.

As shown in Figure 4A, K-Neighbors Classifier, Decision Tree, Random Forest, Feed-Forward Neural Network, Support Vector Machine, Logistic Regression, Deep Learning ANN, and Naive Bayes performed with F1-Scores of 0.891, 0.908, 0.901, 0.904, 0.910, 0.893, 0.908, and 0.881, respectively, with 5-fold cross validation and 0.888, 0.873, 0.882, 0.901, 0.877, 0.899, 0.897, and 0.884 with leave-one-out cross validation (LOOCV). The prediction accuracies and AUC values were also calculated shown in Figure 4B and 4C. Overall, all 8 models performed well with high F1-scores (>88%) and high AUC (>95%) with Support Vector Machine performing the best.

A confusion matrix was plotted for each classification model as shown in Figure 5. The matrices summarize the number of true positives, false positives, true negatives, and false negatives among the reader, writer, and eraser predictions generated by each model. The results from these prediction models are cataloged into the CREWdb.

**Figure 5:**
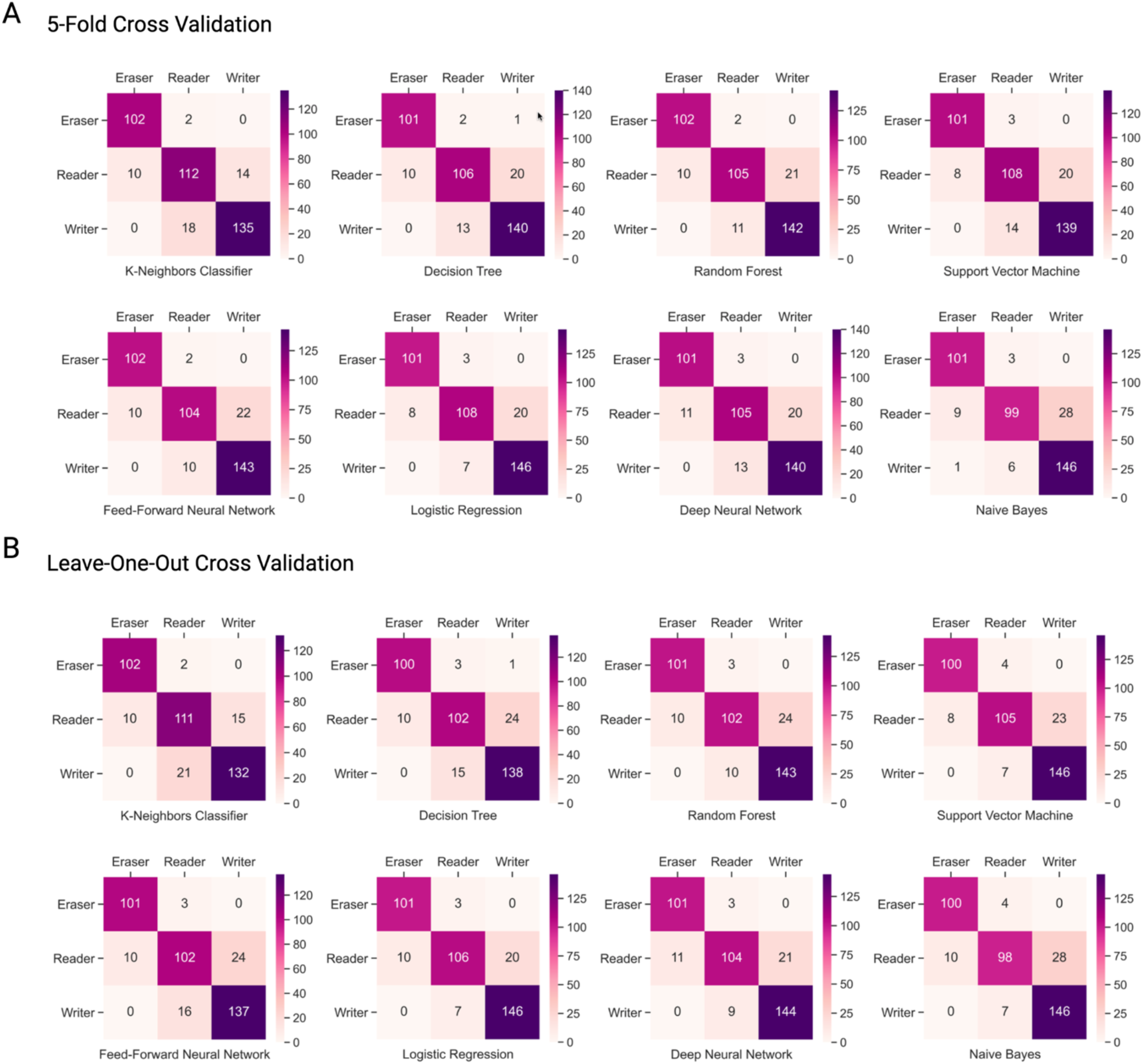
(A) Confusion matrices of models run with five-fold cross-validation. (B) Confusion matrices of models run with leave-one-out cross-validation.

## Web interface and utilities

The web application offers two major utilities. The first utility allows the user to browse chromatin remodeler proteins by specifying diseases and disease-subtypes, to obtain the corresponding chromatin remodeling functional roles and their likelihood of their role membership. The user selects the diseases of interest, their subtypes from the ‘Diseases’ and ‘Subtypes’ dropdowns respectively. The search text box inputs a list of protein names or EntrezIDs with each protein on a new line. On clicking the submit button, the web interface sends the user input as a server request and the result section is populated using the information stored in CREWdb. The output/results is divided into two sections – result tables and knowledge graph representation (Figure 6).

**Figure 6:**
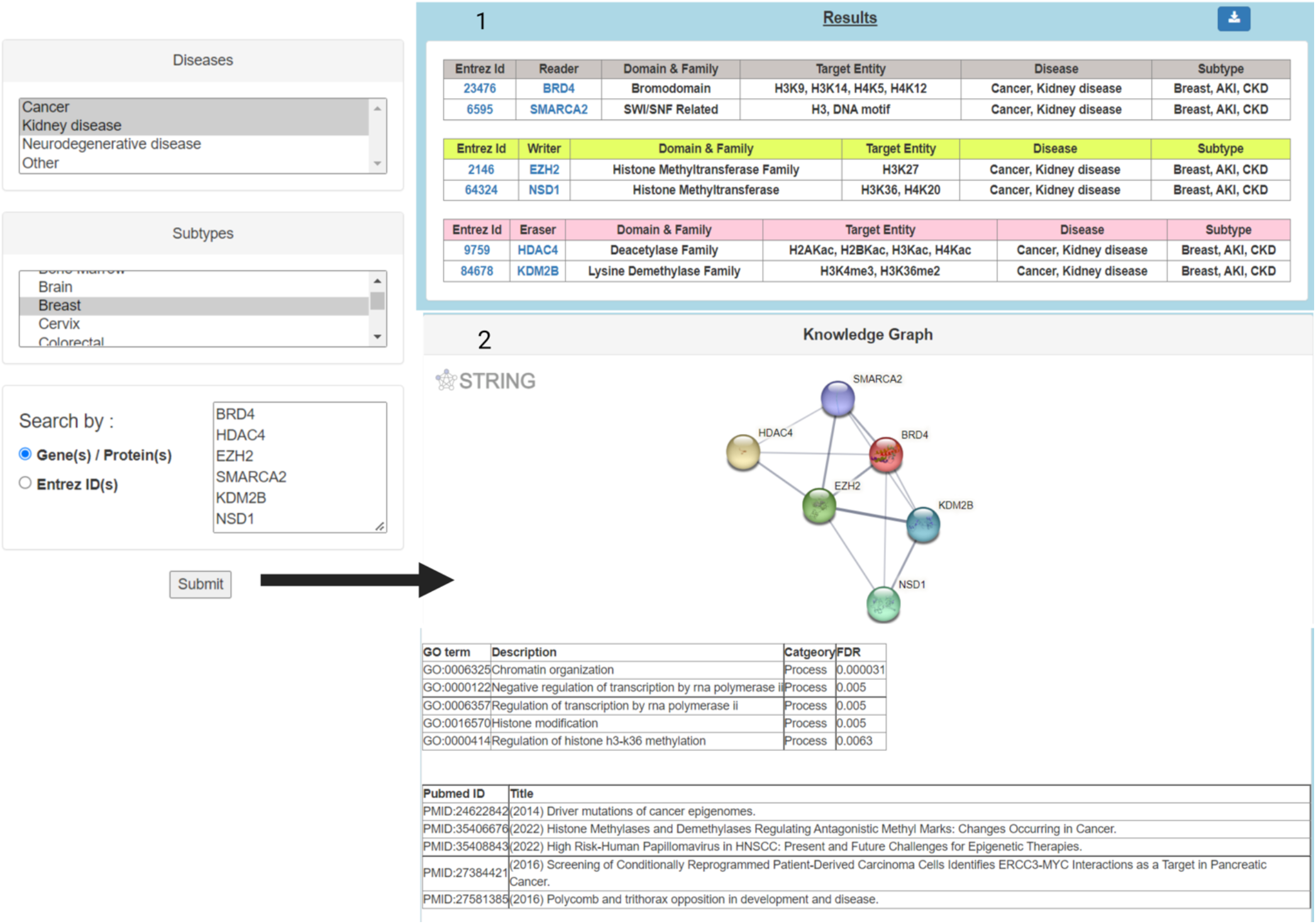
Utility of the CREWdb website. A list of proteins is entered as input in the ‘Search by’ box after selecting desired diseases and subtypes. The output section is divided into two parts – 1) result tables and 2) knowledge graph.

The first utility of the output section - result tables contains a list of tables with each of input proteins classified as reader, writer, or eraser respectively. Each of the role tables is retrieved using the CREWdb tables containing protein information – EntrezID, domain and family name, and percent probabilities for each functional role and their associations with specific histone marks (top part of Figure 6, 7). On clicking the EntrezID cell for a protein in the table, its corresponding GeneCards page opens in a new tab. Similarly, the user can also view the probability/likelihood of the given protein as a reader, writer, and eraser by hovering over the protein name. When a protein is classified as two or more functional types, the probability of the respective type is displayed as “TBD”. On clicking the EntrezID for EZH2, the user is directed to its GeneCards page displaying genomic related information of EZH2 as a writer. A Naïve Bayes machine learning model is used to display the probability of a given protein being a reader, writer, or eraser. The download function allows users to export the results in CSV (comma-separated values) format (bottom part of Figure 7).

**Figure 7:**
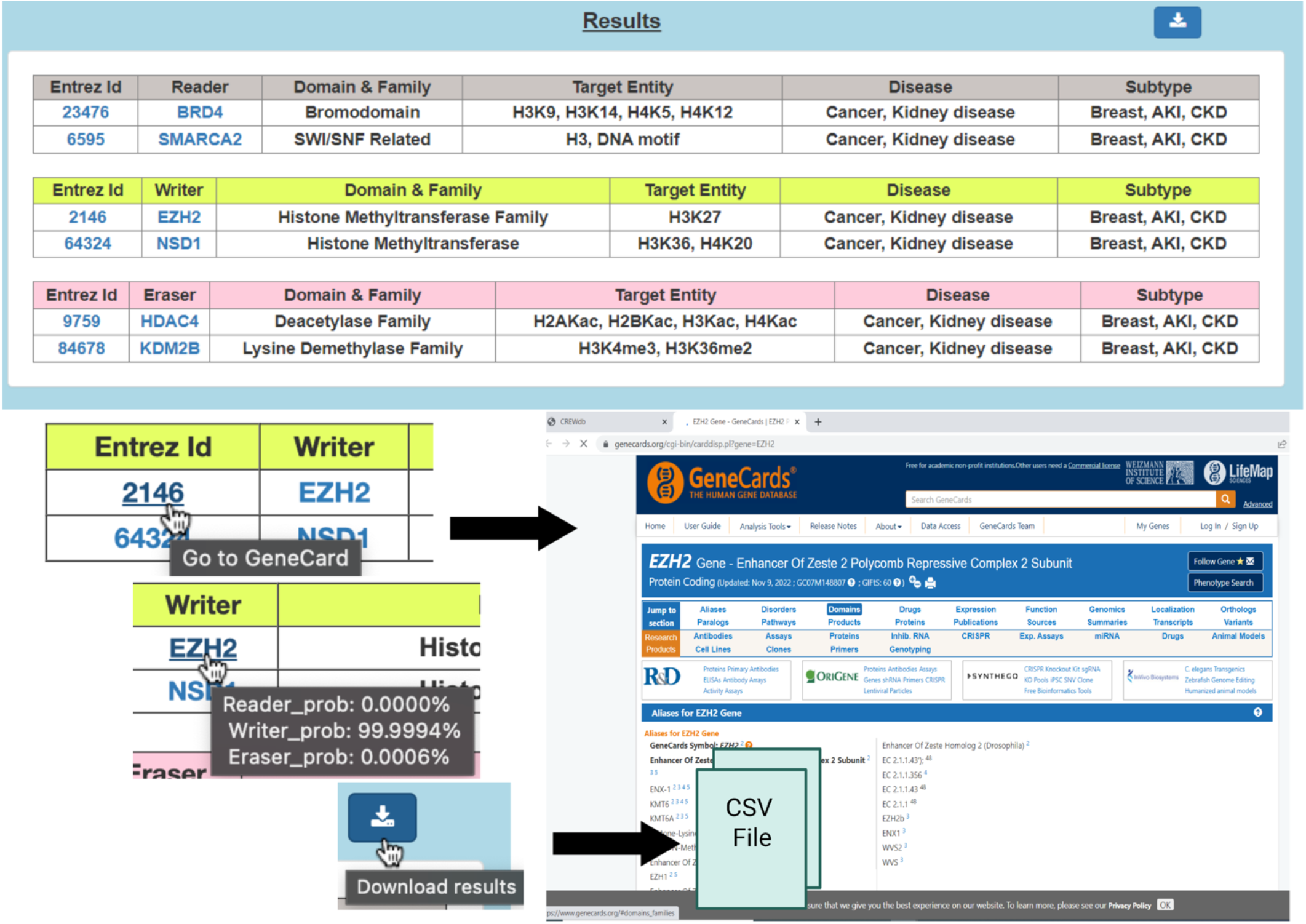
The results table section of the output with utilities to browse GeneCards, view functional probabilities of the chromatin remodelers and export the tables as csv files.

The second utility is a visualization tool that generates the network structure for the protein list based on their functional interactions used to populate the knowledge graph section. The knowledge graph section is the other part of the output that contains the network structure, functional enrichments, and PubMed references for a given protein list. It is populated using StringDb API calls for embedding the interactive protein-protein interaction (PPI) network, obtaining the functional enrichments and corresponding PubMed references. The PPI network captures the interactions among the proteins, such that the strength of the interactions is determined by the thickness of the edge connecting them. By clicking on the protein of interest (for example EZH2), the user can view the AlphaFold protein structure, additional protein information and also browse other resources. On hovering over the protein name, for example, EZH2 in the results table (Figure 8), the knowledge graph section displays the PPI network, functional enrichments, and Pubmed references for EZH2 (Figure 8). All default parameters of StringDB API (species–9606 (human), network flavor-confidence, caller-identity) were used to populate the knowledge graph output section.

**Figure 8:**
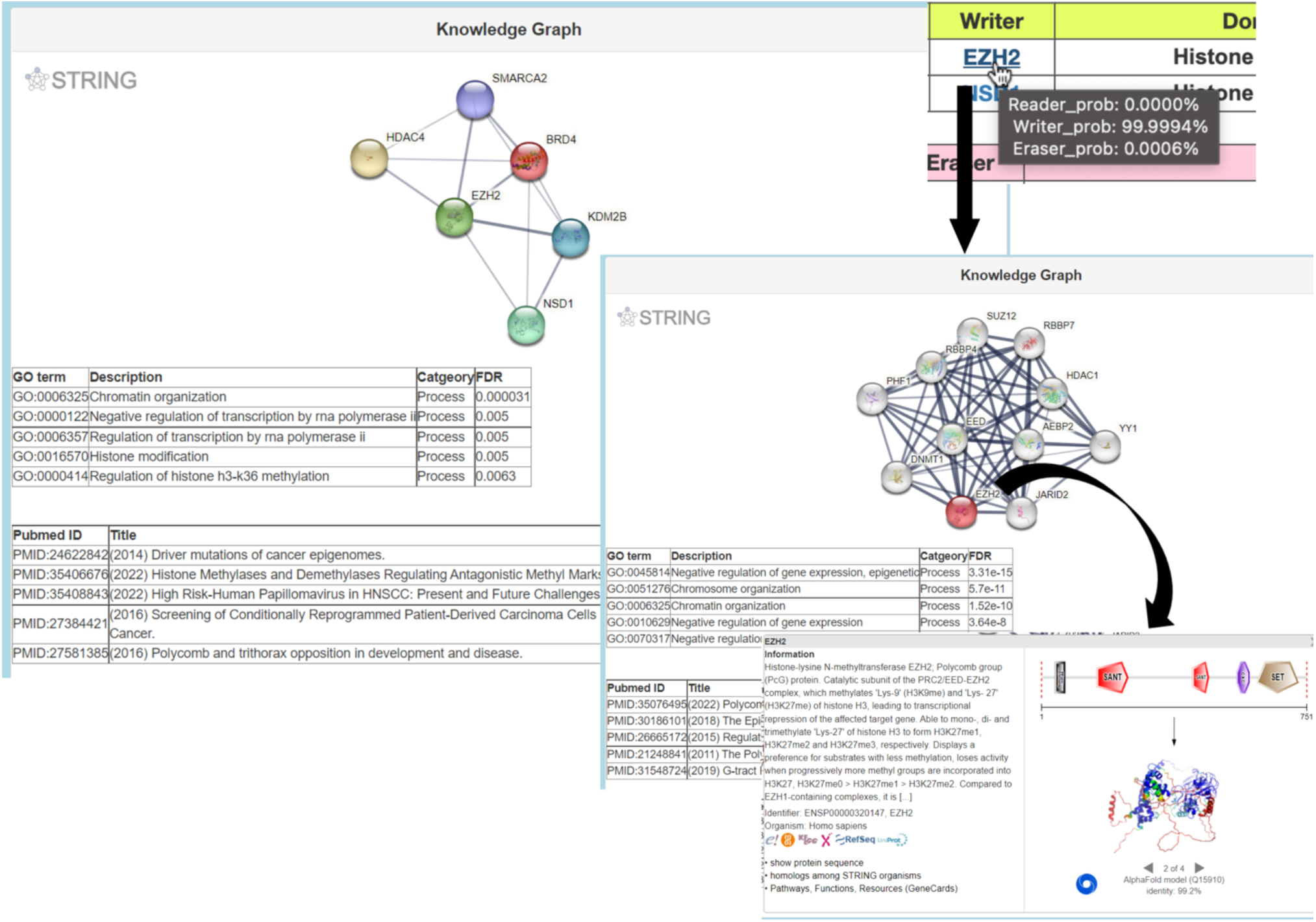
Utility of the knowledge graph capturing, protein-protein interaction network, functional enrichments, and PubMed references for the protein/s in question and their meta information (e.g., protein structure conformation, interactive protein partners, etc).

## Discussion

With the initial data curation of chromatin regulators, we found overlaps between classified chromatin regulators as readers and writers or readers and erasers. Of the 1484 chromatin regulators extracted, 133 proteins were classified as both readers and writers and 67 proteins were classified as both readers and erasers. This significant overlap reduces the clarity in understanding the epigenetic regulation of diseases such as cancer, neurodegenerative diseases, kidney diseases, and other rare diseases.

The lack of clarity could be due to the limited coverage of available validated annotations. Moreover, we believe that the ambiguity in chromatin regulator roles is due to the complexity of their biological and molecular interactions in tissue specific manner. Chromatin regulators have been characterized as enzymes that work as part of a multiprotein complex, which explains why a chromatin regulator protein can act as a reader, writer, or eraser depending on its biochemical context. Given the ambiguity of regulatory roles of known regulators, we included a Bayesian model to provide probabilistic scores to demonstrate the multifaceted nature of a regulator, which can have overlapping roles as a reader, writer, or eraser. For example, proteins such as DNMT3A and SIRT2 have eraser, reader, and writer functionality probabilities 97.3235%, 2.6765%, 0.000% and 1.0350%, 98.9650%, 0.0000%, respectively. DNMT3A is a type of methyltransferase, a protein that transfers methyl groups from one substrate to another. The probabilities derived from the Bayesian model suggest that this protein has functionalities related to readers and erasers. Similarly, the functional probabilities for SIRT2, a protein that removes acetyl and longer chain acyl groups from lysine residues, reflect its potential bifunctionality as a reader and eraser. Furthermore, this model suggests that proteins mainly classified as erasers can also have attributes of those of readers, and vice versa, and illustrates the complexity of chromatin regulator roles.

Generally, knowledge of chromatin regulator functionality is obtained through rigorous wet-lab validations. However, these experiments are often time-consuming and costly. To overcome this barrier, we devised a data-driven approach that extracts biologically relevant features, refines those features, and utilizes them to train machine learning models to classify proteins into their chromatin regulator roles as readers, writers, and erasers.

Considering the machine learning pipeline developed in this study, we specifically trained models on biologically relevant features from the EpiFactors database to predict chromatin regulator roles in cancer. The resulting prediction accuracies indicate that a highly confident feature set yields a good model accuracy. Yet, there are limitations due to the lack of known and validated chromatin regulators, the size of the dataset, and imbalanced class labels. As a result, the interpretation of prediction accuracies may be ambiguous for some regulators.

Given the ambiguity in chromatin regulator roles, it is vital to further understand and identify the unique features of chromatin regulators that contribute to their characterizations as readers, writers, and erasers. By examining the biological significance of each epigenetic feature, we can further identify insightful patterns across extracted data and potential areas of improvement for the machine learning pipeline.

Out of the 9 epigenetic features selected for the machine learning model, *Pfam domain, HGNC gene family description*, and *HGNC gene family tag* are promising features that could be further expanded on for the development of CREWdb. Pfam domains are derived from a database of protein families and utilize a hidden Markov model, which searches for alignments with the greatest matches at conserved sites. Many genes classified within the readers, writers, or erasers categories had similar or the same Pfam domains. Readers were consistently associated with ARID, Bromodomain, and JmjC domains; writers were associated with acetyltransferase, DUF, KRAB, MBD, and methyltransferase domains; and erasers were associated with amino oxidases, ARB2, Cupin, F-box, HDAC, SIR2 domains. Given the association between Pfam domains and chromatin regulator functionality, corresponding Pfam domains could be further inspected to understand the classification of readers, writers, and erasers.

In addition to extrapolating existing features, additional epigenetic information on cofactor interactions and histone modifications or disease-specific data could enhance the predictability of the model. For example, for predicting chromatin regulator roles in cancer, additional features could include RNA expression derived from RNA-Seq data of primary cancer types, correlations between RNA expression and single copy number alteration for specific cancer types; correlations between RNA expression and promoter methylation level for specific cancer types; differential methylation status between tumor and normal samples; proteomics data based on reverse phase protein array for primary tumor tissues and antibody staining level for human normal tissues; correlations between phospho-proteins and histone expressions between tumor and normal samples; and correlations between clinical features such as subtype, overall survival, etc. and RNA expression. In the future we plan to integrate these features to improve the predictability of the chromatin remodeler proteins’ functional roles in epigenetic regulations of diseases in tissue specific manner. We will also utilize text mining techniques to automatically extract chromatin remodeler proteins-feature-disease associations from Pubmed data to further increase the coverage of the CREWdb knowledge base.

## Conclusion

Using our machine learning pipeline, we demonstrate that by utilizing feature enrichment techniques such as feature selection and class rebalancing we can identify highly confident features that lead to high prediction accuracies for chromatin regulator classification. Given the overlap in chromatin regulator roles, we plan to optimize our probabilistic model for each chromatin regulator role as reader, writer, or eraser by integrating disease-specific features. This will further enable our understanding of the complex natures of these regulators’ multifaceted roles in various disease phenotypes. In the future, we plan to extend our CREWdb to a comprehensive knowledge base system by introducing text-mining techniques to extract knowledge from biomedical articles and by integrating other disease knowledge sources. In doing so, we hope to establish the CREWdb as a comprehensive and insightful knowledge source on the functional roles of chromatin regulators across all disease types.

## Acknowledgements

We thank CR2Cancer, TCGA, CCLE, WERAM DB, EpiFactors databases for making data available to us. We also thank Melba Nuzen for helping designing the CREWdb logo.

## Author Contributions

Original concept: S.M.; experimental design: S.M.; implementation: M.N., R.G., M.K., M.S., S.M.; data interpretation: M.N, R.G., M.S., S.M.; manuscript writing: M.N., R.G., S.M.; website development: R.G., S.M.

